# Diminished placental Factor XIIIA1 expression associates with pre-conception antiretroviral treatment and preterm birth in pregnant people living with HIV

**DOI:** 10.1101/2024.10.30.620906

**Authors:** Doty Ojwach, Nicola Annels, Sunny Sunshine, Daniel Simpkin, Justin Duruanyanwu, Tariq Webber, Berenice Alinde, Nadia Ikumi, Michael Zulu, Hlengiwe Madlala, Landon Myer, Thoko Malaba, Marie-Louise Newell, Paolo Campagnolo, Siamon Gordon, Heather Jaspan, Clive M. Gray, Fernando Martinez Estrada

## Abstract

We previously showed a link between maternal vascular malperfusion and pre-term birth (PTB) in pregnant people living with HIV (PPLH) initiating antiretroviral treatment (ART) before pregnancy, indicating poor placental vascularisation. After measuring antenatal plasma angiogenic factors to seek mechanistic insights, low levels of plasma Factor XIIIA1 (FXIIIA1) and vascular-endothelial-growth-factor (VEGF) was significantly associated with PTB at the time closest to delivery (median 34 weeks) in PPLH initiating ART before pregnancy. Knowing that FXIIIA1 is crucial for haemostasis, angiogenesis, implantation and pregnancy maintenance and that expression is found on placental macrophages (Hofbauer cells), we examined placentae at delivery from matching participants who either initiating ART before pregnancy or during gestation. Highest FXIIIA1 expression was on Hofbauer cells but was significantly lower in PTB regardless of HIV infection, but was significantly lower in PPLH in PTB from women who initiated ART before pregnancy. To test the hypothesis that antiretroviral drugs may disrupt vascularisation in the placenta, we used a human umbilical vein endothelial cell (HUVEC) matrigel angiogenesis assay. We identified that addition of pre-treated FXIIIA1-expressing MCSF-and IL-10-induced placenta-like macrophages with physiological concentrations of tenofovir, 3TC, and efavirenz resulted in significantly inhibited angiogenesis; akin to the inhibition observed with titratable concentrations of ZED1301, an inhibitor of FXIIIA1. Overall, an efavirenz-containing ART combination inhibits vasculogenesis without causing toxicity and likely does so through inhibition of a FXIIIA1-mediated-placental macrophage pathway.

## Introduction

Universal antiretroviral therapy (ART) provided to pregnant people living with HIV (PPLH) has mitigated transmission of HIV during pregnancy, but also appears to be associated with complications of adverse birth outcomes, such as preterm delivery and low birth weight (1–5). Such adverse birth outcomes have implications for subsequent growth (6, 7), cognitive developmental defects (8) and increased cardiovascular risks for both the child and the mother (9), and other long-term complications that need ameliorating. Previously we established that pregnant people living with HIV (PPLH) initiating ART during conception, as opposed to those initiating ART after conception, face a two-fold risk of vascular malperfusion (MVM), which in turn significantly associates with preterm birth (3).

We set out in this study to explore the impact of ART initiation on Hofbauer cells, foetal macrophages found in placental villous tissue. Hofbauer cells form a dynamic network that supports spiral artery remodelling, phagocytosis of apoptotic cells, angiogenesis, trophoblast invasion, and promote immune tolerance within the villi (10). They could play a key role in MVM, in its crosstalk with foetal vascular malperfusion (FVM), that includes thrombosis, non-vascularised villi, villous karyorrhexis due to insufficient removal of apoptotic cells, vascular intramural fibrin deposition due to coagulation, stem vessel obliteration, and vascular ectasia (11).

To identify an angiogenic marker in circulating antenatal plasma that could explain the MVM observation, we found that low circulating levels of plasma VEGF and FXIIIA1 were significantly associated with pre-pregnancy ART and pre-term birth. Both VEGF and FXIIIA1 are involved with angiogenesis, with the latter being an enzyme of the transglutaminase family (12–15), participating in the steps required for matrix remodelling and early blood supply to the developing foetus through the placenta. The enzyme is a candidate to control vascular development and permeability during pregnancy in humans. Mice carrying transgenic embryonic trophoblast cells overexpressing or lacking FXIII, show respectively, decreased or increased vascular permeability and altered architecture of the blood vessels at the implantation site, measured in MRI as surface area and fraction blood volume (16). FXIIIA1 is mainly produced by monocytes and tissue macrophages in the body and is exacerbated during alternative activation of macrophages (17, 18). The enzyme has been loosely described as a Hofbauer cell marker (19) and plays a crucial role in angiogenesis by increasing endothelial cell migration, proliferation and survival (16, 20). When looking in placentae, collected at delivery, FXIIIA1 was co-expressed with other macrophage markers such as CSF1R, CD68 and CD163 and appeared a reliable marker of Hofbauer cells. Extending this analysis to the PIMS birth cohort (25), we identified that FXIIIA1 expression and density in placentas from PPLH and specifically when ART was initiated before pregnancy, strongly associated with pre-term birth. We hypothesise that diminished FXIIIA1 expression by placental macrophages is associated with impaired vascularisation, explaining the association between pre-pregnancy ART and MVM. We provide mechanistic evidence that FXIIIA1 is involved with disrupted angiogenesis and explore the hypothesis that ARV drugs are complicit in this effect.

## Results

### Low antenatal plasma levels of VEGF and FXIIIA1 associates with pre-term birth in PPLH who initiated ART before pregnancy

We previously established that pregnant individuals living with HIV (PPLH) who initiated ART before pregnancy, as opposed to during gestation, faced a two-fold increased risk of MVM (3). The association of MVM with preterm birth and low birth weight suggests that a placenta-mediated mechanism likely links long-term ART use with adverse birth outcomes. Taking the same PIMS cohort (3), we selected plasma samples that matched the placentae we collected at delivery and measured 28 analytes at three time points during gestation. The PIMS cohort has been previously described (25), but briefly PPLH were recruited who were receiving ART prior to conception (S, stable group) or who were initiated on ART during gestation (I, initiating group). Figure 1A shows a heatmap of the 28 plasma analytes supervised by different birth outcomes: appropriate for gestational age (AGA), small for gestational age (SGA) and preterm birth (PTB). Examining the heatmap qualitatively, the HIV negative participants delivering pre-term showed an up-regulation of anti-inflammatory cytokines in the 2^nd^ trimester (Figure 1). PPLH giving birth to SGA neonates showed an up-regulation of chemokines, growth factors and pro-inflammatory cytokines at a median of 34 weeks’ gestation (the last time points prior to delivery in the 3^rd^ trimester).

**Figure 1:**
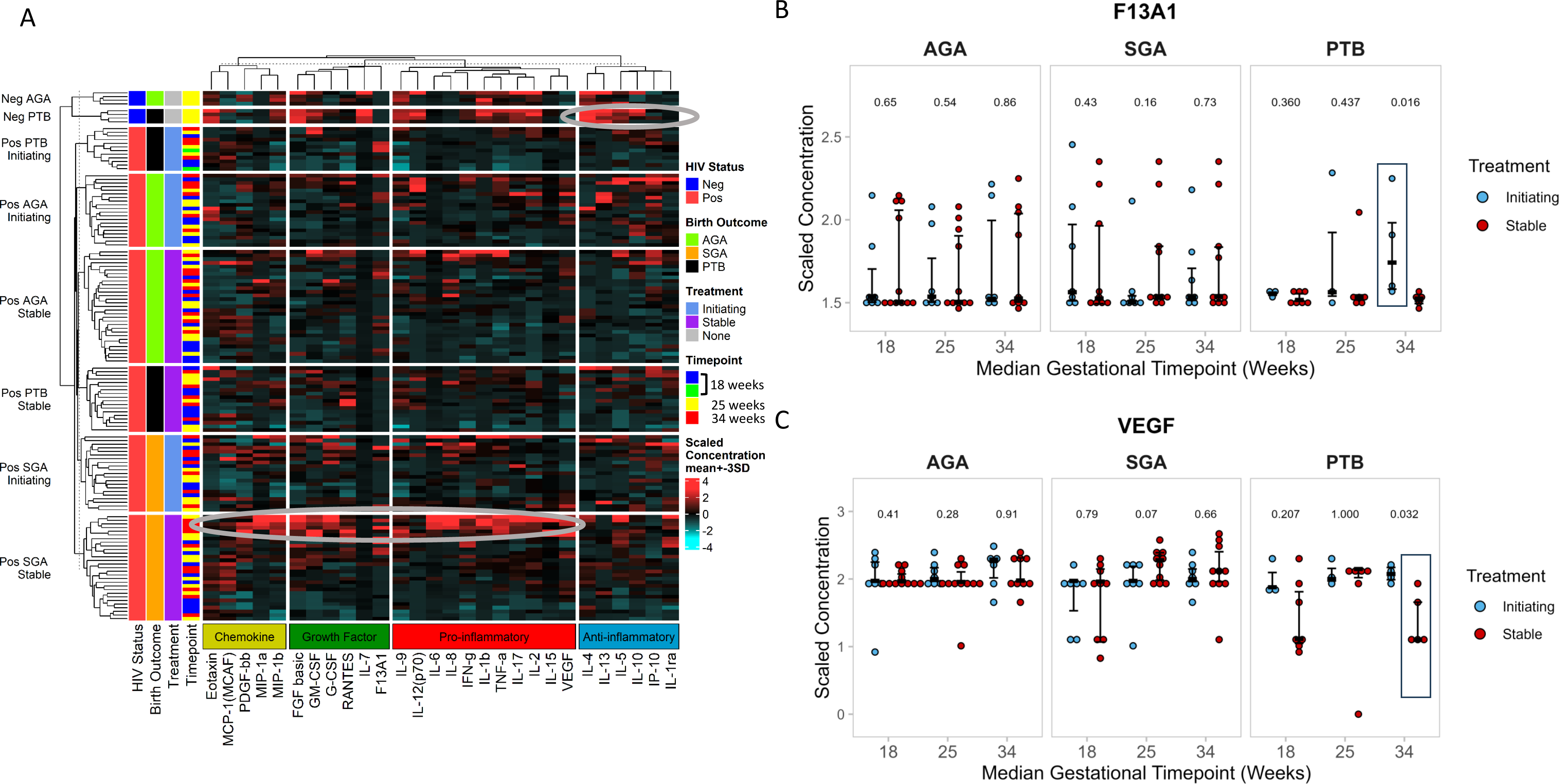
Antenatal expression of 28 plasma analytes during gestation. A. Heat map representation of 28 analytes classified as, chemokines (army green), growth factors (green), pro-inflammatory (red), and anti-inflammatory (blue) markers across PPLH (HIV-positive in red) and uninfected controls (HIV-negative in blue). Participants were further stratified by birth outcomes: Neg (AGA n=4 and PTB n=4) and Pos (AGA n=21, PTB n=11 and SGA n=18). For PPLH, timing of ART is indicated as light blue for those initiating treatment during conception (initiating) and purple for those stable on treatment prior to conception (stable). Grey circles shows the different cytokine profiles for SGA and Neg PTB B. Dot plot comparing plasma FXIIIA1 expression between PPLH groups: those initiating treatment (blue) and those stable on treatment (red) across term-AGA, SGA, and preterm births at median gestational weeks 18, 25, and 34. The black box highlights elevated FXIIIA1 expression in the initiating PTB group at median 34 weeks of gestation. C. Dot plot comparing plasma VEGF expression between PPLH groups: those initiating treatment (blue) and those stable on treatment (red) across term-AGA, SGA, and preterm births at median gestational weeks 18, 25, and 34. The black box highlights depressed VEGF expression in the stable PTB group at median 34 weeks of gestation. Each dot represents one participant at each timepoint, from each plasma analysed. Lines and bars represent the mean and standard deviation, respectively. The Two-tailed P values from Mann-Whitney U test is shown.

When we stratified the 28 cytokines by timing of ART initiation, five cytokines appeared significant at any one of the three antenatal time points comparing stable (S) and initiating (I) groups and birth outcomes: IL-5 and IL-17 at 18 weeks for AGA, IL-17 at 18 weeks for PTB, FXIIIA1 and VEGF at 34 weeks for PTB (Table S1). In AGA at 18 weeks, there was higher levels of IL-17 in initiating vs stable (p=0.023) while the opposite was observed in PTB (p=0.049, Supplementary Figure S1A). Additionally, IL-5 was significantly higher in initiating vs stable in AGA (p=0.0012, Supplementary Figure S1C). Although IP-10 in AGA at 18 weeks did not reach significance (p=0.056), this marker of inflammation was notable at 34 weeks of gestation in initiating vs stable (p=0.036, Supplementary Figure S1B). Of note, Figure 1B shows low levels of FXIIIA1 at median weeks 18, 25 and 34 in PPLH with a PTB outcome receiving stable (pre-pregnancy) ART (Figure 1B). At the later time point closest to delivery in this PTB group (median 34 weeks), circulating FXIIIA1 was significantly elevated (p=0.016) when ART was initiated during pregnancy (initiating group, Figure 1B). Closely allied to FXIIIA1 in function is vascular endothelial growth factor (VEGF) which was lower at the time point closest to delivery (a median of week 34) and with PTB (p=0.032, Figure 1C). However, the depressed VEGF was found only in the stable group (ART prior to pregnancy) which is further depicted with the clustering of PTB initiating and stable groups in the PCA plot, suggesting that VEGF and FXIIIA1 plasma levels may differentiate the PTB group according to initiation of treatment, Supplementary Figure 1D. The direction and length of the vectors for VEGF (closer to PC2, 44.5%) and FXIIIA1(closer to PC1, 55.5%) indicate their contribution to the variance represented by each principal component. PC1 with 55.5% of the variance captures most of the variation, suggesting both FXIIIA1 and VEGF may drive the primary differences in PTB birth outcome and timing of treatment. The PTB stable group (red dots) have both low FXIIIA1 and VEGF expression, while the PTB initiating group (yellow dots) have balanced expression of both with a bias towards elevated FXIIIA1 potentially linking these two cytokines to the PTB outcome in the stable group. Together these data show that plasma markers related to vascularization are depressed in PPLH prior to pre-term birth in those who initiated ART before conception.

Tentative evidence that either plasma concentrations of FXIIIA1 and/or VEGF could be potential predictive markers of pre-term birth is shown when constructing Receiver Operating Characteristic curves. Supplementary Figures 1E-G show marginally significant area-under-the-curves (AUC) for plasma FXIIIA1 (AUC, 0.8; p=0.07) and significant VEGF (AUC, 0.889; 0.019) and combined FXIIIA1+VEGF AUC, 0.8278, p=0.0047), indicating the close association between depressed circulating angiogenic factors and PTB. Collectively, these data suggest an association between pre-conception ART and disruption of pathways related to FXIIIA1 and VEGF production and PTB and potential vascular development.

### Expression of F13A1 gene and FXIIIA protein in the placenta

To seek clarity on how FXIIIA1 may be implicated in MVM and PTB, we investigated its expression within the placenta in two ways. First, an *in silico* analysis from publicly available data (21) was performed and second, was to verify surface protein expression in term placentae matching the plasma analysis shown in Figure 1. Figure 2A shows *in silico* analysis of these datasets and demonstrates the *F13A1* gene co-expression with *CSFR1*, *CD68* and *CD163*. It is known that Hofbauer cells express an array of expressed genes: CD68, CD14, CD64, CD206, FOLR2, LYVE1+, CD163L1, CD36, F13A1, FGF13, LYVE1, MEF2C, SPP1(22, 23) and macrophage colony stimulating factor receptor (CSF1R) (24). The *in silico* analysis revealed that the *F13A1* gene is expressed during first trimester placentae and specifically by Hofbauer cells (Figure 2A). Verification of Hofbauer cell-expression of FXIIIA1 staining in our term placentae is shown in Figure 2B – demonstrating anatomical location of single-stained FXIIIA1, CSFR1, CD163 and CD68 within trophoblasts. When staining our term placentae simultaneously for FXIIIA1, CSFR1, CD68 and CD163, we showed that these macrophage surface markers were variably co-expressed within trophoblasts (Figures 2C, D and E) and demarcated the location of Hofbauer cells. Levels of marker co-expression revealed that FXIIIA1 was majorly co-localised with CSF1R (83.3%), CD68 (80%) and then CD163 (30.8%) (Figure 2D and E) and mainly co-expressed on CD68+CSFR1+ cells (89.2%, Figure 2D). Staining exhibited a diffuse cytoplasmic granular pattern, predominantly in villous tissue and within the trophoblasts (Figures 2B and supplementary Figure 2A). Supplementary Figure 2A shows that FXIIIA1 single staining is dispersed sparingly in the maternal decidua parietalis and basalis, and focused predominantly within the Hofbauer structures in the villous tissue. In the decidua, there were two patterns of FXIIIA1 staining, “striated” exhibiting a dimmer and stellated appearance, yet the cells are less abundant and compressed, attributed to the heightened density of the matrix. Other sections showed staining of cells aggregating in “lumps” some seemed artificial but are a characteristic of transglutaminases in tissue (Supplementary Figure 2B. In DP there was association of FXIIIA1 staining with smaller nuclei.

**Figure 2:**
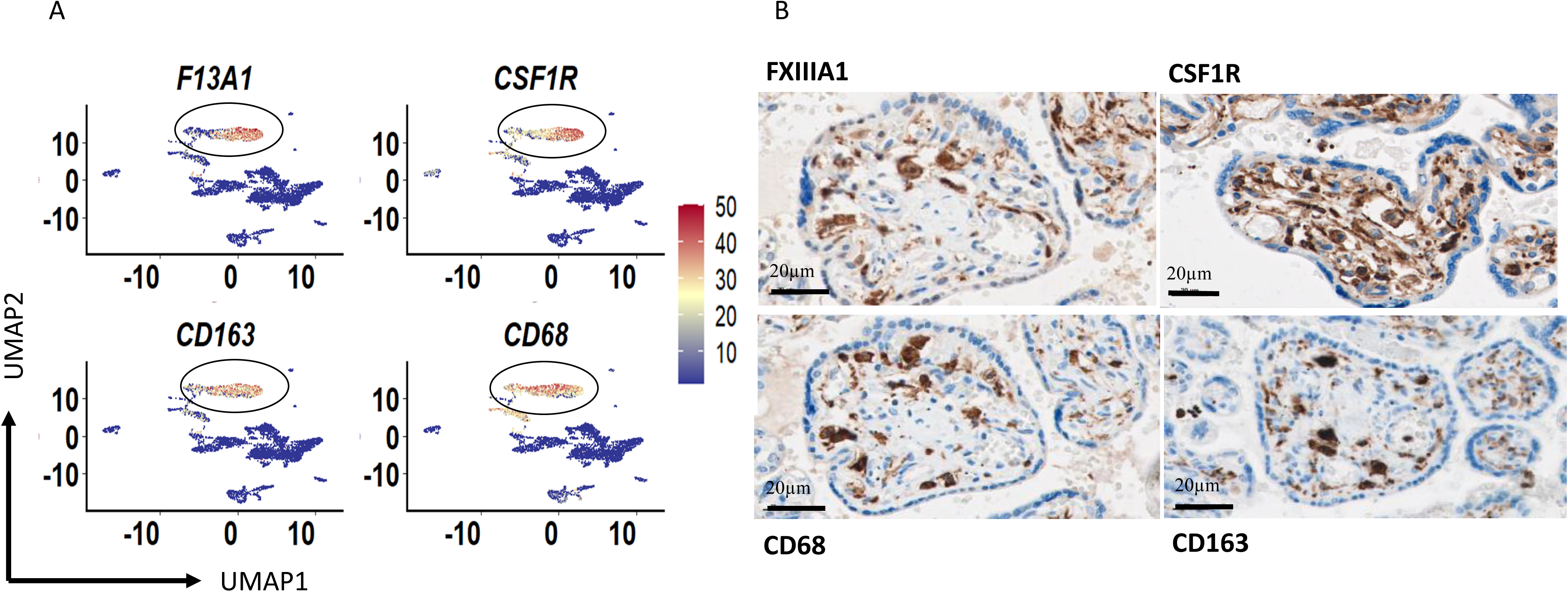
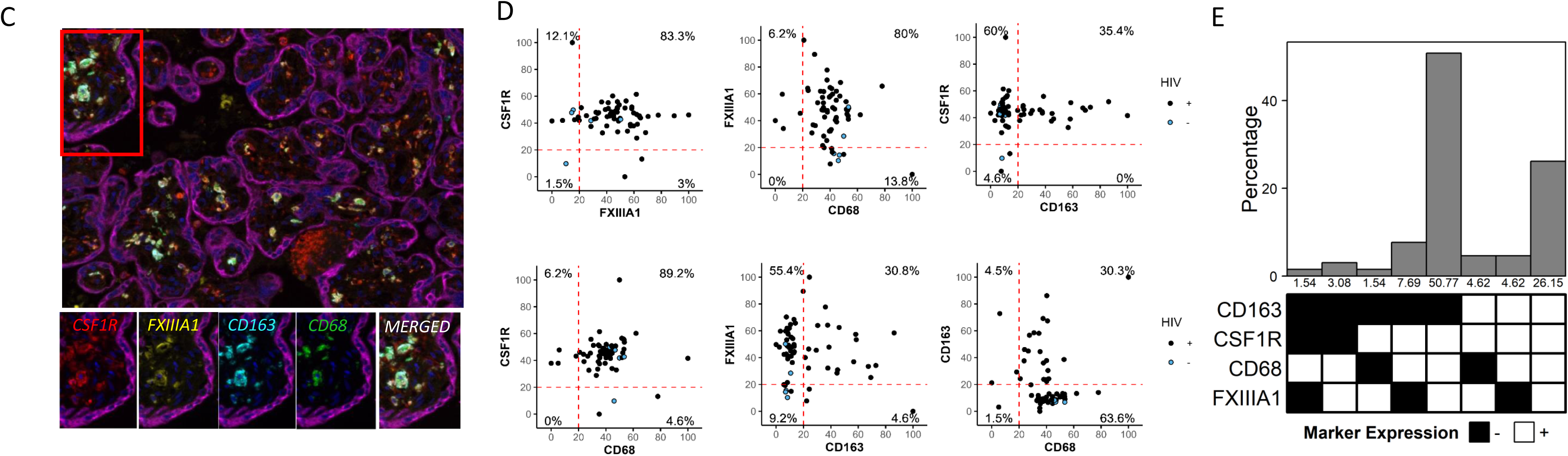
FXIIIA1 expression and other macrophage marker co-expression in Hofbauer cells. A. Uniform Manifold Approximation and Projection (UMAP) analysis of *F13A1* gene expression overlayed with other known macrophage markers *CSF1R*, *CD68* and *CD163*, from single cell data set (21) obtained from 1st trimester placentas. Red to blue denotes high to low expression according to the scale bar. B. Representative images of single marker expression of FXIIIA1, CSF1R, CD68 and CD163 staining of a placental sample collected from a healthy term pregnancy. Scale bar at 20µm. C. Representative of n=11 multiplex immunofluorescence staining image of a placental sample collected from a healthy term pregnancy confirming co-expression of FXIIIA1 Coumarin (yellow) with macrophage markers CSF1R Cy5 (red), CD163 Cy3 (cyan), CD68 FITC (green) in villous tissue. Pan-cytokeratin Alexa 750 (magenta) was used to demarcate the villi and villous trophoblasts. Nuclei was visualised using DAPI. The overlay depicts the merged signals. D. Dot plots showing co-expression of FXIIIA1 with CSF1R (83.3%), CD68 (80%), CD163 (30.8%). Blue and black dots represent co-expression of FXIIIA1 analysed from HIV-and HIV+ placentas. E. Representation of the percentage co-expression of all the 4 various macrophage markers. White and black squares represent expression and no expression

### Low Hofbauer expression of FXIIIA1 in placentae from PPLH and associates with pre-term birth

Quantifying expression of FXIIIA1 in placentae tissue showed lower density expression in the decidua tissues (DP and DB separately) and higher cell density in the villous tissue (Figure 3A). Notably, FXIIIA1 expression was significantly lower in villous tissue from placentae collected from PPLH when compared with term placentae from HIV negative women. When we stratified FXIIIA1 staining in the villous tissue between PPLH initiating ART before conception (S) or during gestation (I), there was no difference in FXIIIA1 expression (Figure 3A).

**Figure 3:**
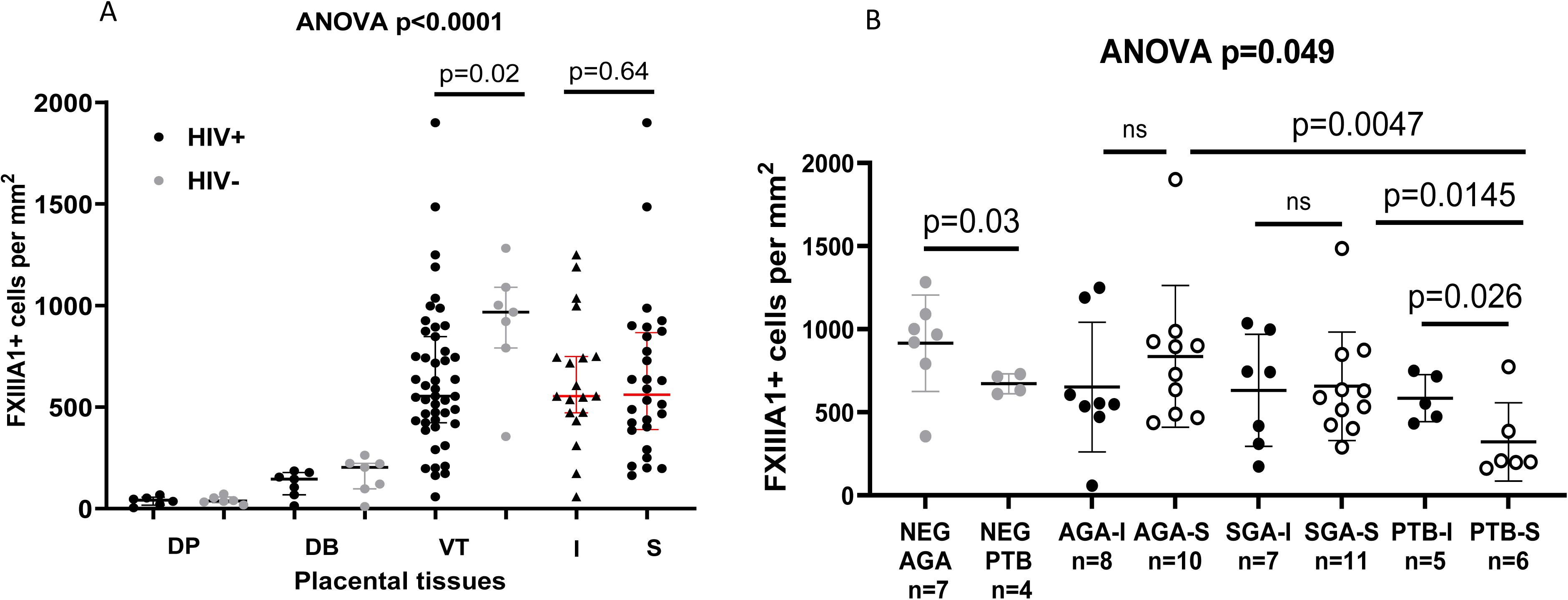
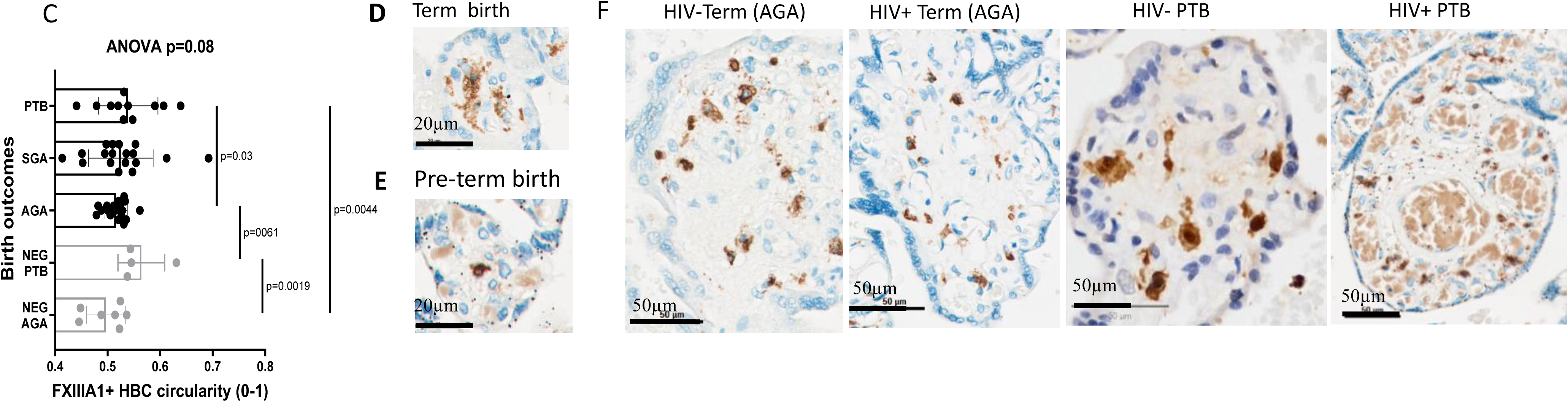
FXIIIA1+ Hofbauer cells distribution in birth outcomes and initiation of antiretroviral therapy Dot plot). A. Dot plot quantifying FXIIIA1+ cells within placental membrane regions: fetal chorionic villous tissue (VT), maternal decidua parietalis (DP), and decidua basalis (DB). Black and gray represent HIV+ and HIV-placenta respectively. The HIV+ group (black) is further categorized by antiretroviral treatment timing: pre-conception (Stable-S, dots) and post-conception (Initiating-I, triangles) B. Dot plot quantifying FXIIIA1+ cell expression patterns and variations in birth outcomes. Black and gray represent HIV+ and HIV-placenta. Filled black circles denotes initiating treatment and open circles stable on treatment. C. Quantitation of morphological feature of Hofbauer cells circularity ANOVA p=0.08 (C) and D. representative placental sections showing hofbauer cell in term (D) and E. pre-term placenta (E). Scale bar 20µm. F. Representative placental sections used for quantification in B, are displayed from term appropriate gestational age (AGA) and preterm birth (PTB) cases, including both HIV-negative and HIV-positive individuals. Each dot represents mean ± SD of 5 regions of interest from each placenta analysed. Lines and bars represent the mean and standard deviation, respectively. The overall repeated measures ANOVA p value is shown as well as the the Two-tailed P values from Mann-Whitney U test is shown.

We next examined FXIIIA1 expression in placentae from HIV negative women at term (range: 856-1046 cell/mm^2^) and pre-term (range: 627-686 cell/mm^2^) and identified that there was significantly lower expression in PTB (p=0.03, Figure 3B). Upon measuring FXIIIA1 expression using the samples from the PIMS cohort (25), there was no difference between the density of these cells between AGA and SGA, regardless of ART timing (Figure 3B). The density of FXIIIA1 in placentae from PTB initiating ART during pregnancy was no different from SGA or AGA births or PTB from HIV negative women. Notably, there was a significantly lower FXIIIA1 expression in PTB from women who initiated ART prior to pregnancy (S group).

When examining all samples, regardless of gestational age at delivery, there was a significant positive correlation between FXIIIA1+ cell density and infant birth weight (Supplementary Figure 3A), and marginal positive correlation with gestational weeks (Supplementary Figure 3B) suggesting that FXIIIA1 cell density is proportional to how well the neonate develops *in utero*.

### Morphological differences in FXIIIA1-expressing Hofbauer cells between birth outcomes

To have some idea of differences in macrophage function, we measured the perimeter, area, and circularity of Hofbauer cells in our single DAB-stained samples, with a value of 1.0 signifying a perfect circle and decreasing values indicating progressively elongated shapes. These metrics are proxies for irregularities in cell adhesion, spreading, interaction with other cells, motility, proliferation, and differentiation, including changes in cell size due to different environmental milieu (26–28). We observed a hierarchy of macrophage size as measured by FXIIIA1 staining circularity according to birth outcomes: NEG-PTB>PTB>SGA>AGA> with the least circular cells found in HIV negative (NEG)-AGA controls (Figure 3C). Representative images in Figures 3D and E show high circularity at term (Figure 3D) and low circularity in pre-term (Figure 3E). The opposite was observed in the area and perimeter covered by the cells where the hierarchy was: AGA>SGA>PTB>NEG-PTB, with HIV negative (NEG)-AGA controls having the largest surface area and perimeter, Supplementary Figures 3C-D. To highlight the relationship between area and circularity, we used the 2D kernel estimation density plots to visualize the distribution of Hofbauer cells according to birth outcomes, Supplementary Figures 3E. This suggests that not only are the numbers of FXIIIA1+ cells lower in PTB, but Hofbauer cells are smaller, occupy less space with less FXIIIA1 expression. Figure 3F show representative images comparing FXIIIA1 distribution within trophoblasts to highlight differences between a term placenta from a healthy control (HIV-Term) vs a placenta from PPLH (HIV+ Term) vs pre-term birth in controls versus a PPLH on pre-conception ART. In the latter, there was an increased presence of bloated capillary vessels, indicative of villous immaturity and potential compensatory mechanisms for restricted vascularization, possibly linked to maternal vascular malperfusion.

Collectively, these data show that FXIIIA1 expression by Hofbauer cells is depressed when the birth outcome is pre-term. However, the finding that pre-conception ART is the main driver of low FXIIIA1 expression (Figure 3B), would indicate that pre-conception antiretroviral drug exposure is disrupting the expression of this Hofbauer-bound enzyme. Based on these data, along with our earlier finding of MVM in the same cohort, we hypothesised that pre-conception ART was disrupting vascular development through FXIIIA1 on placental macrophages.

### FXIIIA1 supports vascularisation in an HUVEC angiogenesis assay

To test our hypothesis, we used the conventional HUVEC matrigel model to first assess the role of FXIIIA1 in promoting angiogenesis. Upon activation of endothelial cell receptors by proangiogenic signals such as basic fibroblast growth factor (bFGF), vascular endothelial growth factor (VEGF), platelet-derived growth factor (PDGF), and epidermal growth factor (EGF), endothelial cells release proteases that degrade the basement membrane (29–31). This process allows the endothelial cells to proliferate and migrate, leading to the formation of new sprouts. Figure 4A shows the details of the experimental design, where the initial aim was to confirm that exogenous FXIIIA1 could support angiogenesis. Figure 4B shows images of branch formation when FXIIIA1 was added and then could be inhibited by the addition of ZED 1301 (a FXIIIA1 inhibitor). Figure 4C (condition 3) shows that addition of 5ug/ml of active FXIIIA1 induced a significant increase in branching over and above growth media alone (condition 2), but not necessarily tube thickness (Figure 4D, conditions 2 and 3). Titration of the FXIIIA1 inhibitor ZED1301 concentrations counteracted the effect of adding exogenous FXIIIA1 by reducing both the number of branches and tube thickness in a stepwise manner. Together, these data demonstrate the ability of FXIIIA1 to promote vasculature development.

**Figure 4:**
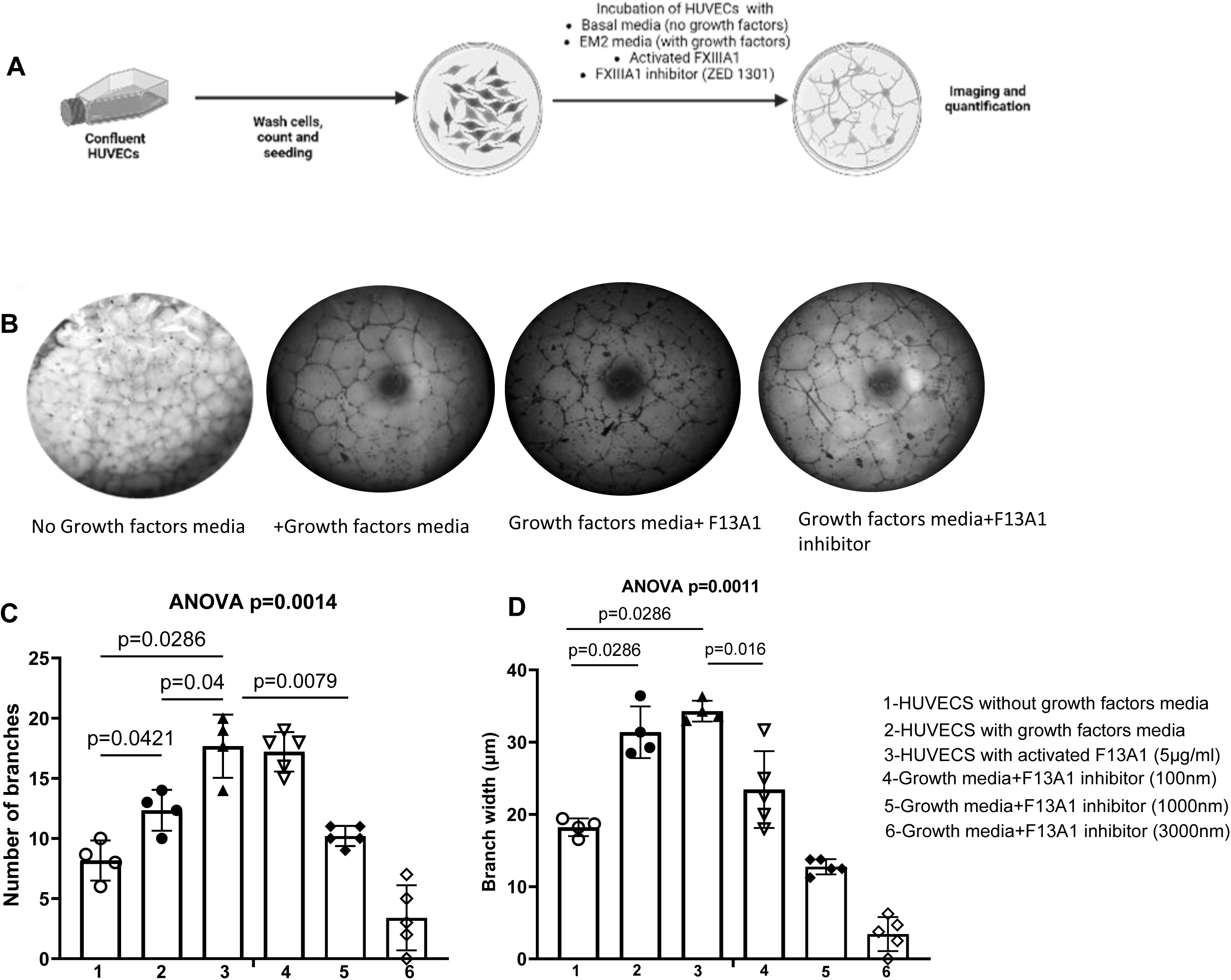
FXIIIA1 supports branch formation and branch thickness in HUVEC angiogenesis assay. A. Experimental design of the HUVEC assay. B. Grayscale images at 10X magnification showing branch formation under various culture conditions: with and without growth factors media, activated F13A1, and its inhibitor ZED1301. C. Quantification of branch formation (ANOVA p=0.0014) and D. Branch width (ANOVA p=0.0011) under different culture conditions: 1) HUVECs without growth factors media, 2) with growth factors media, 3) with activated F13A1 (5 µg/mL), 4) with growth factors media and ZED1301 (100 nM), 5) ZED1301 (1000 nM), and 6) ZED1301 (3000 nM). The design included n=3 experimental repeats, performed in duplicates. The overall repeated measures ANOVA p value is shown as well as the unpaired T-test p values from two-way comparisons. No growth factors media is the basal media for HUVECS. Growth factors media is the EGM2 which contains supplements to support growth of endothelial cells.

### Antiretroviral drug disruption of HUVEC angiogenesis

To test if pre-conception ART was disrupting vascular development through FXIIIA1 on placental macrophages, we generated placenta-like macrophages by stimulating primary human monocytes with M-CSF and IL-10 *in vitro* as previously described (32). Macrophages polarized with 20 ng/mL of IL-10 showed the highest expression of FXIIIA1, as confirmed by flow cytometry (Figures 5A and B) and western blots using a FXIIIA1 monoclonal antibody, detecting FXIIIA1 at 80 kD (Figure 5C). When we tested the ability of macrophages stimulated with either IL-4, MCSF, IL-10 or lipids (as control) in the HUVEC matrigel assay, only the IL-10 cultured macrophages could support branch and tubule width formation over and above the other culture conditions Figures 5D, E and F. We thus regarded these FXIIIA1-expressing cells to be analogous to placental macrophages (32) and used these cells in the HUVEC model to test the impact of ARV drugs.

**Figure 5.**
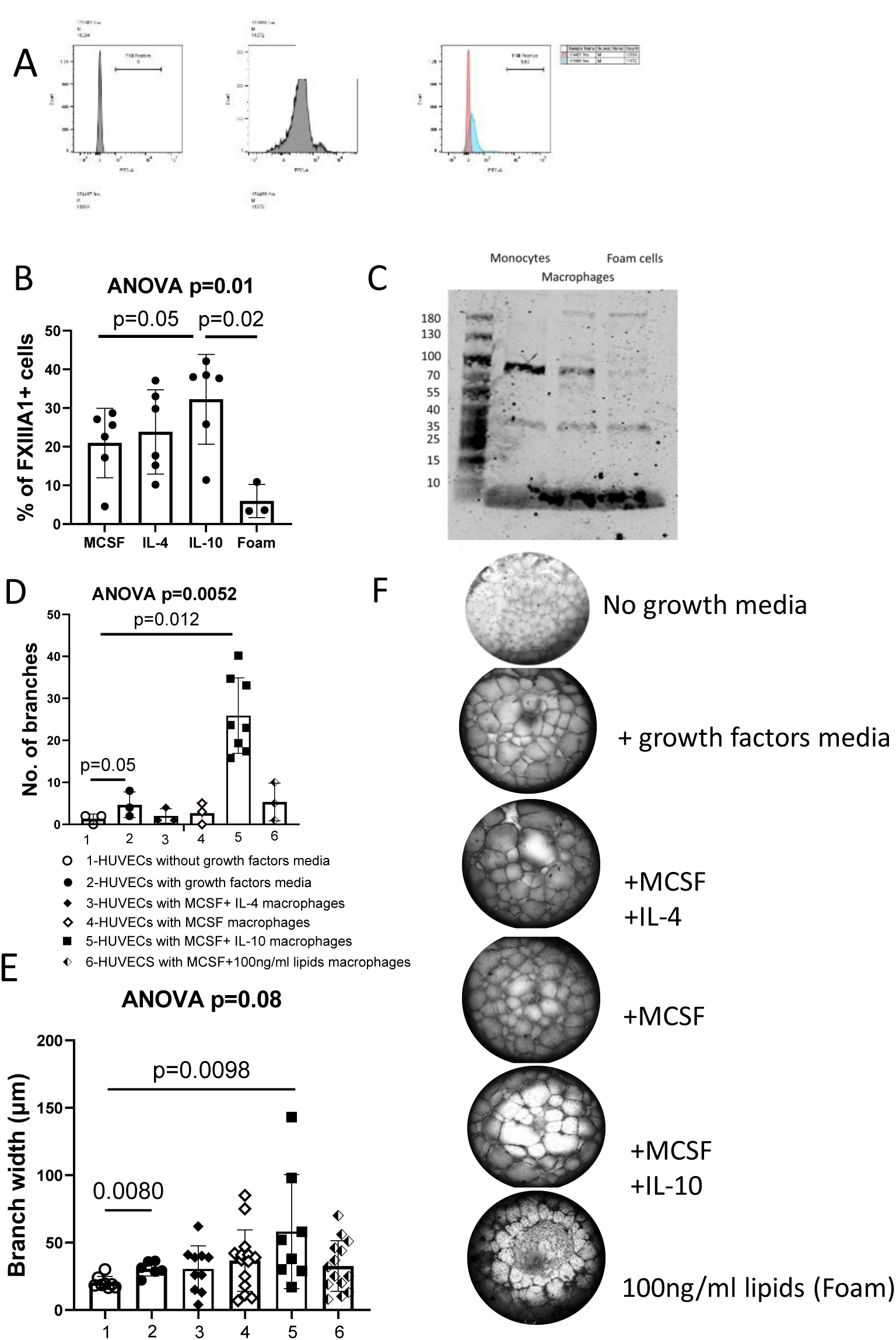
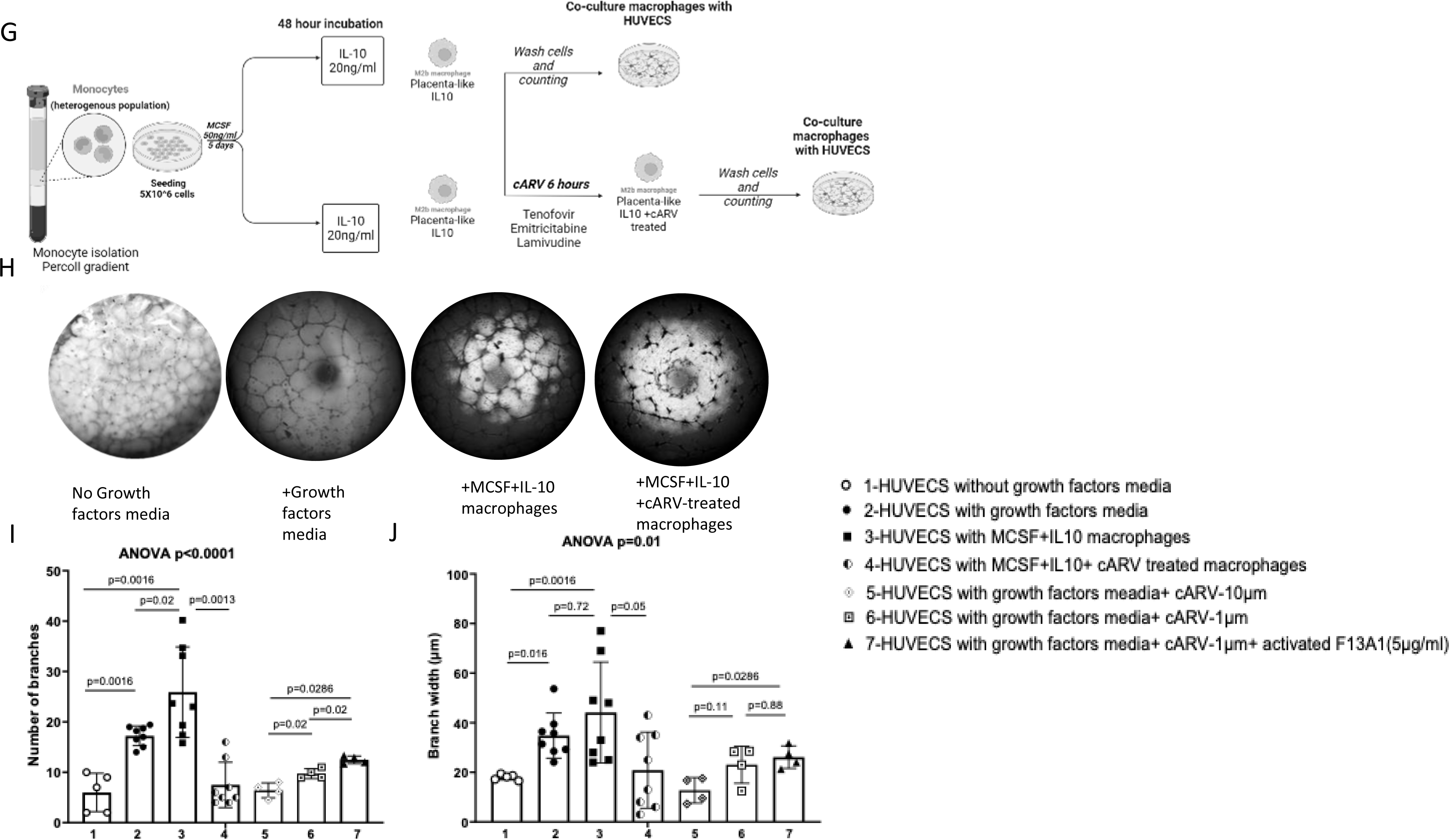
FXIIIA1 Expression in Polarized Macrophages and Disruption of HUVEC Angiogenesis by Placental-like Macrophages Treated with Combined Antiretroviral Drugs. A. Histogram plot showing the gating for negative and positive FXIIIA1 expression using FlowJo software, with pink representing the negative threshold and blue showing positive expression. B. Dot plot showing the percentage of FXIIIA1 expression in macrophages polarized with 20 ng/mL of MCSF, IL-4, IL-10 and 100ng/ml lipids (Foam cells). C. Western blot confirming FXIIIA1 expression at 80 kD in monocytes, macrophages, and foam cells D. Quantification of branch numbers and E. Branch width across culture conditions: 1) HUVECs without growth factors media, 2) with growth factors media, 3) with MCSF+IL-4 macrophages, 4) with MCSF macrophages, 5) with MCSF+IL-10 macrophages, and 6) with MCSF+lipid macrophages (100 ng/mL). F. Grayscale images at 10X magnification showing branch formation in HUVECS with the addition of growth factors media, IL-4, MCSF, IL-10 and foam macrophages. G. Experimental setup for macrophage polarisation, treatment and co-culture with the HUVEC assay. H. Grayscale images at 10X magnification showing branch formation with addition of growth factors media, MCSF+IL10 macrophages and cARV-treated macrophages. I. Quantification of branches (ANOVA p<0.0001) and J. Branch width (ANOVA p=0.01) under different culture conditions: 1) HUVECs without growth factors media, 2) with growth factors media, 3) with MCSF+IL-10 macrophages, 4) with MCSF+IL-10+cARV-treated macrophages, 5) with growth factors media+cARV (10 μM), 6) with growth factors media+cARV (1 μM), and 7) with growth factors media+cARV (1 μM) + activated FXIIIA1 (5 μg/mL). The design included n=4 experimental repeats, performed in duplicates. The overall repeated measures ANOVA p value is shown as well as the unpaired T-test p values from two-way comparisons. No growth factors media is the basal media for HUVECS. Growth factors media is the EGM2 which contains supplements to support growth of endothelial cells.

Figure 5G shows the experimental design, where these polarized placental-like macrophages were pre-treated with non-toxic concentrations of a combination of tenofovir, emtricitabine and lamivudine (supplementary Figure 4) and then added to the HUVEC assay. Importantly, adding the IL-10-polarized placental-like macrophages to the HUVEC assay could significantly (p=0.0016) support the formation of angiogenesis in the absence of supplementary growth media (Figures 5H and I (condition 3) and J (condition 3)). When these macrophages were pre-treated with the combination of tenofovir, emtricitabine and lamivudine, mimicking the regimen of the PIMS cohort (25), both the numbers of branches and tubule width (Figures 5I and J, condition 4) were disrupted in the HUVEC model (p=0.0013 and p=0.05 respectively). When the combined ARV drugs were added directly to the HUVEC assay, in the absence of placenta-like macrophages, there was a similar disruption of angiogenesis (Figure 5I and J, conditions 5 and 6). This effect could be partially rescued by the addition of exogenous activated FXIIIA1 (Figure 5I and J, condition 7).

## Discussion

Inflammatory markers and cytokine profiles vary across gestational time points (33), birth outcomes (34, 35), and the timing of ART initiation in PPLH. By analyzing plasma from multiple time points during gestation, our data suggest that timing of ART initiation significantly influences the immune and vascular profile in pregnant people living with HIV (PPLH), with distinct patterns emerging based on ART timing and birth outcomes. For those initiating ART before pregnancy, a reduced expression of angiogenic factors FXIIIA1 and VEGF was observed close to delivery in cases resulting in preterm birth (PTB). The reduction of these plasma analytes suggest compromised vascularization, with FXIIIA1 implicating a placental component which could contribute to adverse birth outcomes. Conversely, individuals who initiated ART during pregnancy exhibited elevated levels of FXIIIA1 nearer to delivery in PTB cases, suggesting possible immune activation influenced by the proximity of ART initiation. Additionally, cytokine profiles varied by ART status, IL-5 and IL-17 show divergent trends depending on whether ART was initiated pre-pregnancy or during gestation. Among HIV-negative individuals who delivered pre-term, there was an increase in anti-inflammatory cytokines early in the second trimester, suggesting an adaptive mechanism distinct from that seen in PPLH, who showed heightened pro-inflammatory and growth factor signalling closer to delivery in SGA cases. Together, these findings underscore that ART initiated before pregnancy may result in a sustained but potentially maladaptive vascular response, whereas ART during pregnancy seems to trigger a more acute immune shift. Further insights could be critical in understanding ART’s impacts on pregnancy and may guide personalized ART strategies to improve birth outcomes in PPLH.

FXIIIA1 expression by Hofbauer cells is thought to contribute to the maturation of chorionic villi (36). The presence of placental FXIIIA1+ cells was identified as early as the 5^th^ week of pregnancy (19), characterized by developing phagocytic vacuoles and intensified FXIIIA1 staining in tandem with placental growth until approximately 7 weeks of gestation (19). In our study, staining for FXIIIA1 in term placentas from healthy pregnancies, showed that FXIIIA1 expression coincided with an array of known markers for Hofbauer cells and is therefore a useful marker for foetal macrophages. The connection between tissue transglutaminase (TGM2), FXIII, and maternal vascular development during embryo-maternal interactions (19, 20) emphasizes the importance of linking FXIII to angiogenesis during implantation, as reported in murine models (16). Our previous study demonstrated an association between heightened risk of maternal vascular malperfusion (MVM) in PPLH receiving ART before conception, significantly associating with preterm birth and low birth weight, but not when ART initiation occurred during gestation (3).

As FXIIIA1+ cell densities were significantly lower in placentas from both HIV uninfected pregnant people and PPLH with PTB, this represents an important link between adverse birth outcomes and the role of FXIIIA1 expression in the placenta. Our exploration of birth outcomes revealed notable patterns, with FXIIIA1+ cell density being higher in term HIV negative and HIV positive appropriate-for-gestational-age (AGA) compared to preterm births. The lower number of these cells, regardless of maternal HIV infection, and its association with PTB underscores the role of Hofbauer cells and/or FXIIIA1 function in maintaining term pregnancies. Moreover, the timing of ART initiation further influenced FXIIIA1 expression, as placentas from those initiating ART before pregnancy exhibited the lowest FXIIIA1+ expression and PTB. The central question is whether long-term ART gives rise to poor vascularization due to macrophage dysfunction, leading to poor placental development.

The mechanism by which pre-conception ART may be contributing to pre-term birth is supported with our findings of placental-like macrophages, which express FXIIIA1 after polarization with IL-10, play a crucial role in supporting angiogenesis in the placental environment (30), a process potentially disrupted by antiretroviral drugs (ARVs). The observed ability of IL-10-stimulated macrophages to significantly support tubule formation in the HUVEC assay aligns with previous studies demonstrating the angiogenic potential of diverse macrophages (37). Notably, the reduction in angiogenic branching and tubule width following exposure to a combination of tenofovir, emtricitabine, and lamivudine mirrors prior reports linking ART to placental vascular alterations and adverse pregnancy outcomes, such as MVM, SGA and preterm birth (4, 5). This disruption is indicative of ART’s effect on placental macrophage function and subsequent vascular development, which has been highlighted in studies assessing maternal vascular malperfusion and placental insufficiency associated with ART exposure (3). Moreover, the partial rescue of angiogenesis with exogenous FXIIIA1 along with inhibition of angiogenesis with a FXIIIA1-inhibitor, underscores the central role of this transglutaminase enzyme in maintaining placental vascular integrity, as previously reported in contexts of wound healing and vascular remodelling (30). These results, therefore, provide valuable insights into the mechanisms through which ART may impair placental vascular development, with significant implications for improving therapeutic strategies in pregnancies affected by HIV. Further exploration of the downstream effects of FXIIIA1 on placental macrophage function and vascular development could inform strategies to mitigate the adverse effects of ART on placental health.

Complications such as preeclampsia and intrauterine growth restriction (IUGR) have been linked to disturbances in early uterine blood supply (38), and impaired trophoblast cell invasion of the placental bed spiral arterioles (39). The potential role of FXIIIA1 in these scenarios warrants consideration. In comparing plasma FXIII, the IUGR group showed consistent FXIII levels, while the normal pregnancy group experienced significant reductions (40). This suggests that the decline in plasma FXIII levels in normal pregnancies could be attributed to maternal FXIII transfer through the placenta to the foetus.

Taken together, our data suggest that PPLH receiving pre-conception ART are at a higher risk of PTB and that a possible mechanism involves the combined effects of ARV drugs on angiogenesis through FXIIIA1 and Hofbauer cells. If verified with further studies, our findings advocate that adjunctive therapy be provided to mitigate the hypo-vascular effects of pre-conception ART on the placenta during pregnancy.

## Materials and Methods

### Placenta Cohort

This study analysed samples from the Prematurity Immunology in HIV-Infected Women and their Infants Study (PIMS) cohort (25). Briefly, PIMS, was a prospective cohort, aimed to explore the correlation between ART and adverse birth outcomes among women with HIV receiving antenatal care services at Gugulethu Midwife Obstetric Unit (MOU), Cape Town, South Africa (25). Eligible participants encompassed pregnant women aged ≥18 years, and recruitment took place between April 2015 and October 2016 (25). Enrolled people, consisted of those who had initiated ART prior to pregnancy (stable group) or at 14-15 weeks during gestation (initiating group). The ART regimens in this cohort consisted predominantly of tenofovir (TDF), lamivudine (3TC) or emtricitabine (FTC) and efavirenz (EFV)(25). In this present study, matching antenatal plasma samples and placentas with known birth outcomes were analysed: preterm delivery (PTB n=11, less than 37 weeks’ gestation), small for gestational age (SGA, n=18, birth weight less than the 10^th^ percentile), and appropriate for gestational age (AGA, n=22, birth weight ≥25th percentile). Controls included seven AGA term placentas from people without HIV infection recruited via a parallel observational cohort in a neighbouring MOU (Khayelitsha Site B).

Maternal and new-born characteristics by HIV status are presented in Table S1. There were no significant differences in age of the people enrolled. Both groups had median gestational ages of 38 weeks (PPLH, n=51) and 40 weeks (HIV-negative, n=7) respectively. PPLH exhibited earlier deliveries (38 weeks) than HIV-negative (40 weeks, P < 0.0166), with lower placental weight (p<0.05) and new born birth weight (P<0.0189, n=51).

### FXIIIA1 plasma quantification and cytokine profiling

Plasma from 18, 28, 34 weeks prior to delivery were used from the PPLH group for detection of coagulation Factor XIIIA1 polypeptide using a bead-based assay (Cloud-Clone Corp, Katy, USA) and 27 human cytokines, chemokines, and growth factor biomarkers (Bio-Rad, Hercules, California) following the manufacturer’s instructions. The 27 analytes were eotaxin, fibroblast growth factor (FGF)-basic, granulocyte colony-stimulating factor (G-CSF), granulocyte-macrophage colony-stimulating factor (GM-CSF), interleukin-1β (IL-1β), IL-RA, IL-2, IL-4, IL-5, IL-6, IL-7, IL-8, IL-9, IL-10,IL-12, IL-13,IL-15, IL-17, IFN-γ, IFN-γ-inducible protein 10 (IP-10), macrophage inflammatory protein 1α (MIP)-1α (CCL3), MIP-1β (CCL4), monocyte chemotactic protein 1 (MCP-1), platelet-derived growth factor (PDGF)-BB; regulated on activation normal T cell expressed and secreted (RANTES or CCL5), tumor necrosis factor alpha (TNF-α), and vascular endothelial growth factor (VEGF). The concentration of each analyte was determined from a standard curve derived from a human recombinant protein and therefore represents an estimate of the analyte concentration in each sample.

The analyte measurements were performed using the recommended concentrations of reagents and serum, but in half the volume for the three gestational time point collections. This modification has been previously utilized (33). The multiplex plate was read using a Bio-Plex 200 suspension array system (Bio-Rad).

### Placenta tissue collection and processing

Placentas were collected within 6 hours after birth and a portion dissected into decidua parietalis (DP), basalis (DB), and villous tissue (VT). Following fixation in 10% formalin and paraffin embedding, tissue sections (3-5μm) underwent immunohistochemical staining. Complete placentas were fixed in 10% buffered formalin for histopathology, evaluated according to the Amsterdam Placental Workshop Group consensus statement (41). We previously conducted a comprehensive analysis of these placental components, membranes, and the umbilical cord (42).

### Immunohistochemistry

To visualize Hofbauer cells within placental tissue sections, we stained for CD163, CD68, CSF1R and FXIIIA1 markers. After deparaffinization in xylene, and gradual rehydration in a series of ethanol, endogenous peroxidase was blocked by incubating the sections with 0.3% hydrogen peroxide in methanol. Antigen retrieval of placental sections was performed in either a sodium citrate solution (10 mM, pH 6.0) or EDTA (pH 9.0) buffer for a duration of 12 minutes. The tissues were blocked for 15 minutes using 2.5% horse serum (VECTASTAIN PK7200) before incubation overnight with diluted primary antibodies CD163 (Cell Signalling 93498S, 1:400), CD68 (Agilent M0876, 1:400), CSF1R (Abcam 228180, 1:1), and F13A1 (Abcam 1834, 1:300) in 1% BSA/PBS. A biotinylated universal secondary antibody, horse anti-mouse/rabbit (VECTASTAIN PK7200), was applied for 30 minutes at room temperature. HRP coupled Avidin/Streptavidin-biotin complex (ABC Avidin biotin-complex) kit Elite, was used to amplify the signal and visualised with 3,3’-diaminobenzindine (DAB) (Vector SK4100) and counter-stained with haematoxylin (Vector H-3404). Placental sections were dehydrated and enveloped in permanent mounting medium (VectaMount H-5000) before scanning using the PhenoImager (Akoya Biosciences).

### Immunofluorescence staining

Multiplex Immunofluorescence (mIF) staining was done in the Akoya Biosciences Phenoptics platform to validate colocalization of FXIIIA1 as a Hofbauer cell marker with other established macrophage markers. Slides were dewaxed, rehydrated and fixed with normal buffered formalin (NBF) for 20 minutes. The optimised single IHC antibodies were multiplexed in a tyramide signal amplification (TSA^®^) system including pan-cytokeratin and spectral DAPI in the following order: F13A1, opal 480 (1:150), CD68, opal 520 (1:100), CD163, opal 570 (1:100), CSF1R, opal 650 (1:150); Pan-cytokeratin (1:300)-1 hours, Dig (1:100)-10 minutes, opal 780 (1:25)-1hour. For each cycle of staining, the appropriate heat induced epitope retrieval for all the targets was performed using the appropriate buffer (AR6 or AR9) for 15 minutes followed by incubation with Opal Polymer HRP Ms + Rb (Akoya Biosciences). Then, the samples were incubated with opal fluorophore-conjugated tyramide signal amplification (Akoya Biosciences). The slides were rinsed with a TBST wash buffer after each step. After tyramide signal amplification deposition, the slides were again subjected to heat-induced epitope retrieval to strip the tissue-bound primary and secondary antibody complexes before further labelling. These steps were repeated until the samples were labelled with all five markers and spectral DAPI (Akoya Biosciences). Finally, the slides were mounted in Fluoromount-G (Invitrogen). Single stain and auto fluorescence (AF) controls were used for creating an acquisition protocol in InForm software (Akoya Biosciences).

### Microscopy, image scanning and processing

For Microscopy scanning and imaging, the PhenoImager multispectral imaging system (Akoya Biosciences) was employed followed by unmixing and quantification. Raw images were processed and unmixed using InForm software. The batch processing of the mIHC images was done using a custom-made Image-J script. Briefly, following colour deconvolution, background subtraction and median filtering, the appropriate threshold was determined using the Otsu filter. Analyze Particles was then used to detect and count DAB-stained cells. For all the quantification analysis of cell area, perimeter, circularity and density, five random regions of interest per image were considered.

### Antiretroviral drugs

Drugs were sourced from Thermo Fisher: Tenofovir (catalog no. A0416618), Emtricitabine (FTC, catalog no. A0439315), Lamivudine (3TC, catalog no. A0430523), Efavirenz (EFV, catalog no. 466620050), and Dolutegravir (DTG, catalog no. C4672). The drug combinations used were as follows: Combination 1 – Tenofovir, FTC (Emtricitabine), and EFV (Efavirenz); Combination 2 – Tenofovir, 3TC (Lamivudine), and EFV; Combination 3 – Tenofovir, 3TC, and DTG (Dolutegravir); and Combination 4 – Tenofovir, FTC, and DTG. Dimethyl sulfoxide (DMSO) was used as the vehicle for EFV and DTG, while water was used for Tenofovir, FTC, and 3TC.

The concentrations used were 0.014 μM Tenofovir (43), 0.42 μM FTC (44), 1.262 μM EFV(45), 0.27 μM 3TC (46), and 0.602 μM DTG (47). In the drug combinations, the final concentration was used at 1 μM within the physiological dose and 10 μM which was below the IC_50_ threshold.

### Monocyte isolation, macrophage differentiation and polarization

Human monocytes were isolated from healthy donor buffy coats using a two-step gradient centrifugation process as previously described (24). Monocytes (approximately 1.5 × 10⁶) were seeded into a Lumox dish and differentiated into macrophages in Optimem media supplemented with 10% FBS and 20 ng/mL M-CSF for 5 days at 37 °C in a 5% CO₂ incubator. To induce placental-like phenotypes, macrophages were incubated for an additional 48 hours in Optimem with 10% FBS, supplemented with 20 ng/mL IL-10. As controls, macrophages were treated with 20 ng/mL M-CSF, 100 ng/mL lipids, or 20 ng/mL IL-4. On day 8, the cells were treated with cART, consisting of various combinations of 2 NRTIs and an NNRTI, or a vehicle control, for 24 hours. After treatment, the cells were washed, counted, and used in cell-surface immunostaining and co-culture assays with HUVECs at a ratio of 1:4.

### Cell surface immunostaining and flow cytometry

Macrophages were collected by gentle scraping, washed with Dulbecco’s phosphate-buffered saline (DPBS), and incubated with 1 % BSA Human TruStain FcX™ (BioLegend) for 20 minutes. This was followed by incubation with fluorochrome-conjugated antibodies for specific markers for 30 minutes at room temperature, protected from light. The following antibodies, purchased from BioLegend (San Diego, CA, USA), were used: Zombie NIR live/dead fixable cell stain, CD68-PerCP Cy5.5, F13A1-FITC, TGM2-PE, CSF2R-APC, CD16-BV510, and CD33-BV605. Samples were acquired using a triple-laser BD FACS Celesta (BD Bioscience). Data were plotted and quantified using geometric mean fluorescence intensity (MFI) with FlowJo software version 10 (TreeStar, Ashland, Oregon).

### Western blot assay

Cells (5x10⁶) were lysed using radio immunoprecipitation assay (RIPA) buffer (Thermo Fisher, catalog no. 89900) according to the manufacturer’s instructions for subsequent bicinchoninic acid (BCA) assay (catalog no. 23227). Briefly, the cells were incubated with lysis buffer on ice for 30 minutes, followed by 50% pulse sonication for 30 seconds. The lysates were then centrifuged at 140,000 g for 15 minutes at 4°C. Protein concentration in the supernatant was measured at 562 nm (OD562) using a spectrophotometer. For FXIIIA1 detection, 30 µg of protein was separated by SDS-PAGE and transferred to nitrocellulose membranes (Thermo Fisher, LC2001). After blocking with 5% non-fat milk, the membranes were incubated with primary antibodies: F13A1 (1/1000, Abcam 1834) and GAPDH (6C5, Abcam 8245), following the manufacturer’s protocols. Membranes were imaged using the LICOR Odyssey CLx imaging system.

### Human umbilical vein endothelial cell (HUVEC) Matrigel angiogenesis assay

HUVECs (passages 3-5) were cultured for 72 hours in endothelial cell growth medium (EM2, PromoCell) as previously described (29). Briefly, confluent HUVECs were trypsinized by adding 1 ml trypsin-EDTA, and after 3 minutes of incubation at 37°C in a 5% CO₂ incubator, the trypsin was neutralized with EM2 medium. The cells were counted and resuspended to a concentration of 20-25,000 cells in 250 µl of various culture conditions, which were then plated onto solidified Matrigel in a pre-chilled 96-well plate. The plate and tips were pre-chilled on ice for 30 minutes before adding 40 µl of Matrigel to the wells, which was allowed to solidify at 37°C for at least 30 minutes. To visualize and image the tube networks, 4% paraformaldehyde was added to each well and incubated for 2 minutes at room temperature. Wells were then washed twice with PBS, stained with eosin and hematoxylin, washed again with DPBS, and finally filled with 1 ml fresh PBS to prevent the Matrigel from drying out. The co-culture assay was imaged using a Cytation 5 and EVOS inverted microscope. Quantification of the tube networks was performed using ImageJ software with the Angiogenesis Analyzer plugin (48).

### Data Analysis

To enhance the normality of the residuals, cytokine concentrations were log-transformed unless otherwise indicated. The distribution of continuous data was confirmed by Shapiro–Wilk. The continuous clinical and demographic data of the participants were presented as median and interquartile range (IQR) and compared between, HIV status (HIV+ and HIV-) and delivery (term and preterm) women using the Mann–Whitney U-test. The log-transformed cytokine concentration values were compared between the groups by Welch’s t-test and Mann–Whitney (compare ranks) tests when comparing two groups or Kruskal–Wallis tests for more than two independent groups. Categorical variables and frequencies were compared between the groups by Chi-squared and Fisher’s exact tests. Correlation between FXIIIA1 and clinical variables was determined by Spearman’s correlation coefficient (ρ, rho). We utilised the area under the receiver operating characteristic (AUC) curve to determine the predictive capacity of FXIIIA1 and VEGF for PTB. Probability (p) values < 0.05 were considered statistically significant.

All data were analysed using GraphPad Prism 10.0.2. Data visualizations were created with the ggplot2 package in R, version 4.0.2 R (49) 4.0.2 (50). P-values were adjusted for multiple comparisons using the Benjamini-Hochberg procedure(51).

### Study approval

Ethical clearance was obtained from the Human Research Ethics Committee of the University of Cape Town (reference 739/2014 and reference 263/2015), the University of Southampton Faculty of Medicine Ethics Committee (Reference 12542 PIMS), and the Stellenbosch University Health Research Ethics Committee (Reference N21/02/018). All participants provided written informed consent.

## Data availability

No new codes were generated in the generation of the data

## Author contribution

The design of the PIMS study was conceived by MLN, LM and TM. Experimental design was conceived and planned by F.M., C.G., S.G., D.O., and N.A., D.O., S.S., and N.A. carried out the experiments. Image and data analysis were executed by D.O., N.A., T.W., C.G., and F.M. Sample preparation involved contributions from D.O., N.I., M.Z., B.A., and H.M. D.O., N.A., T.W S.G., F.M., T.M., L.M., C.G., M.L.N and H.J. contributed to the interpretation of the results. D.O. drafted the manuscript with edits and input from LM, MLN, C.G., S.G., and F.M. Overall direction and project supervision was done by F.M. and C.G. All authors provided critical feedback, contributing to research, analysis, and manuscript development.

## Acknowledgements

We thank all the study participants and all the support staff of the PIMS study. Thanks also to expert technical laboratory support from Nicole Prins, Portia Manngo, Fuad Waja and Lizette Fick.

The research reported in this manuscript received funding support from the Eunice Kennedy Shriver National Institute of Child Health & Human Development, part of the National Institutes of Health, through Award Numbers R01 HD080385 (PIMS Prematurity Immunology in HIV-Infected Women and their Infants Study) and R01 HD102050 (MELABO Mechanisms leading to adverse birth outcomes in pregnant women living with HIV). D.O was supported through the African Research Excellence Fund (AREF) [grant # AREF-325-OJWA-F-C0913O], the South African Medical Research Council under a Self-Initiated Research [grant # RP10094], the South Africa National Research Foundation [grant # PSTD2204285217], the Africa-Oxford (AFOX)-visiting fellowship and the Company of Biologists Travelling Fellowship [grant #JCSTF23081194].

---

**Figure S1.**
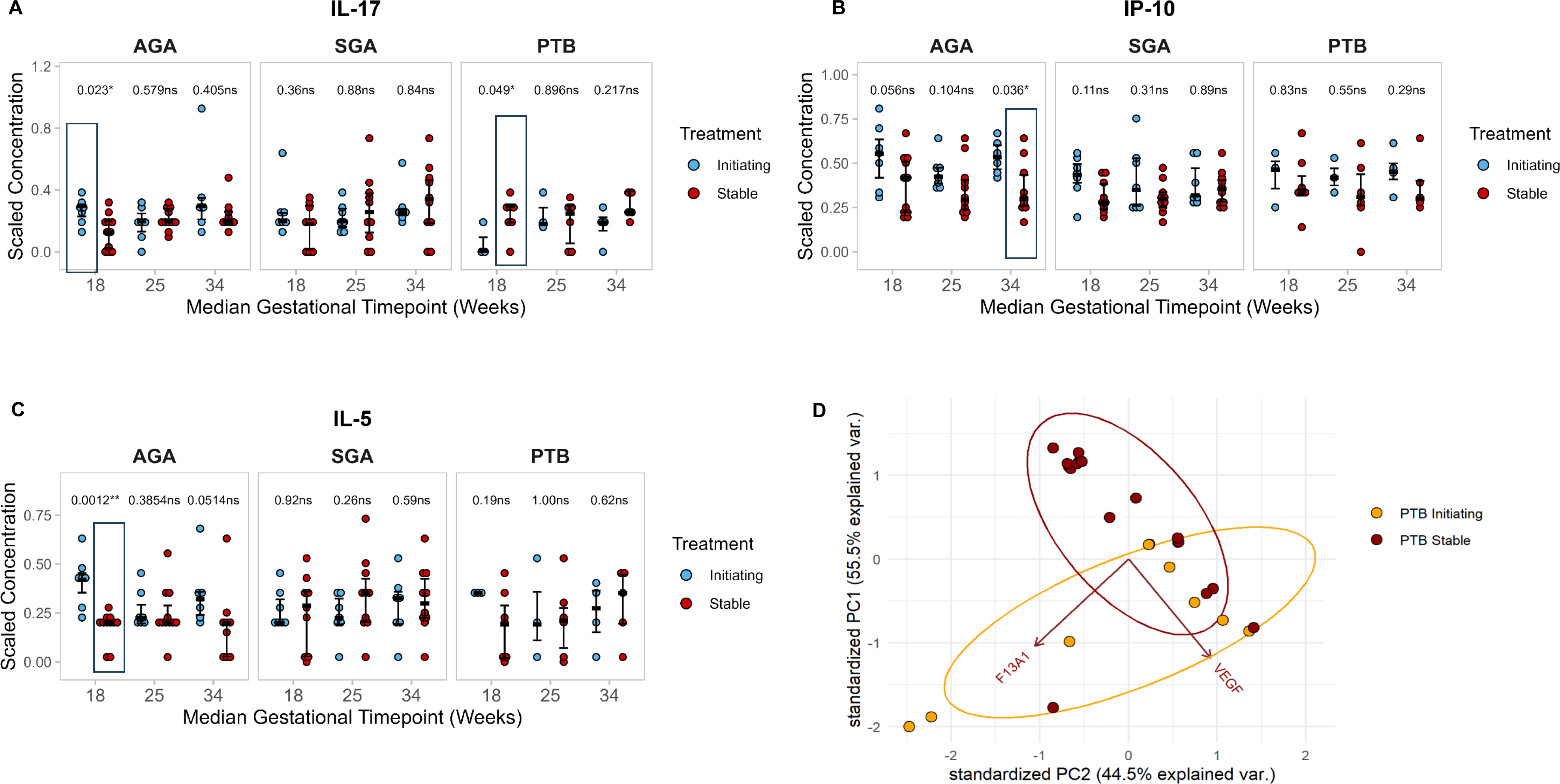
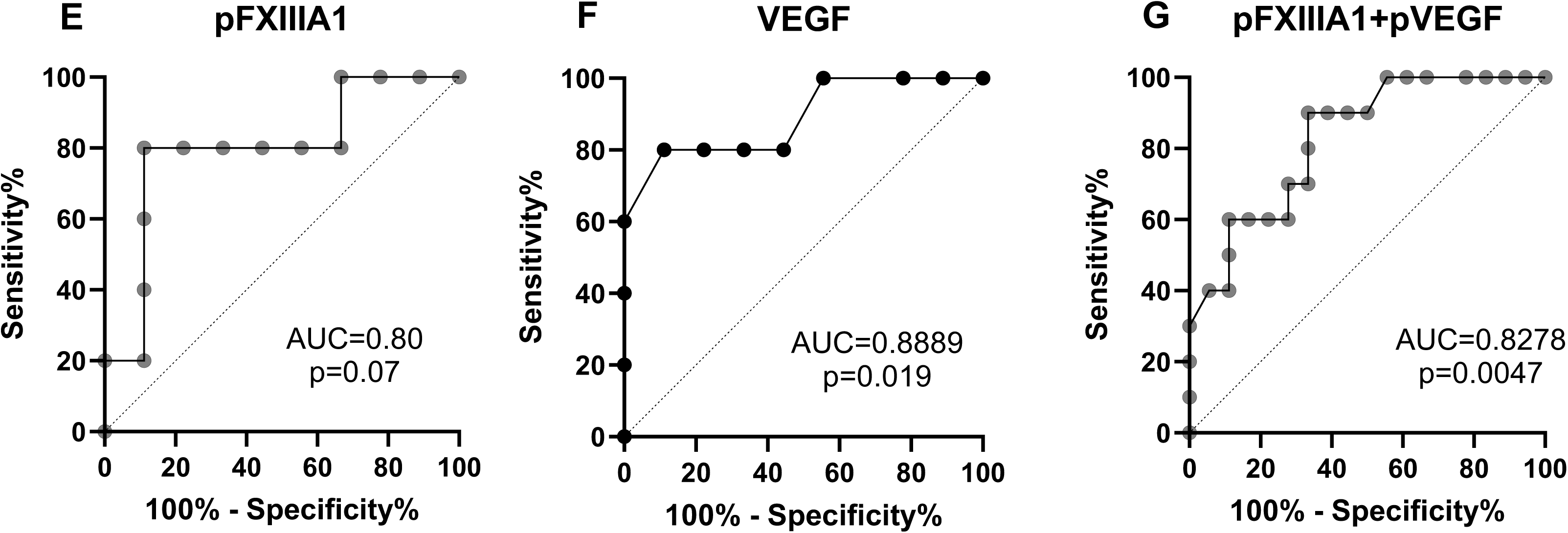
Association of IL-17, IP-10, IL-5, VEGF and FXIIIA1 in PPLH initiating ART with AGA and PTB birth outcomes. A. Dot plot comparing plasma IL-17 expression between PPLH groups: those initiating treatment (blue) and those stable on treatment (red) across term-AGA, SGA, and preterm births at median gestational weeks 18, 25, and 34. The black box highlights elevated VEGF expression in the initiating AGA group at median 18 weeks of gestation and elevated VEGF expression in the stable PTB group at median 18 weeks of gestation B. Dot plot comparing plasma IP-10 expression between PPLH groups: those initiating treatment (blue) and those stable on treatment (red) across term-AGA, SGA, and preterm births at median gestational weeks 18, 25, and 34. The black box highlights depressed IP-10 expression in the stable AGA group at median 34 weeks of gestation C. Dot plot comparing plasma IL-5 expression between PPLH groups: those initiating treatment (blue) and those stable on treatment (red) across term-AGA, SGA, and preterm births at median gestational weeks 18, 25, and 34. The black box highlights depressed IL-5 expression in the stable AGA group at median 18 weeks of gestation D. Principal component analysis of plasma VEGF and FXIIIA1 showing clusters of PPLH preterm birth participants, grouped by those initiating treatment (orange) and stable on treatment (red). Clustering is based on FXIIIA1 (PC1, 55.5%) and VEGF (PC2, 44.5%) concentrations, following batch correction and scaling. E. Plasma FXIIIA1’s role as a predictive indicator of pre-term births in women living with HIV initiating ART before conception AUC=0.80 and p=0.07 F. Plasma VEGF’s role as a predictive indicator of pre-term births in women living with HIV initiating ART before conception AUC 0.89 and p=0.019 G. Combined plasma FXIIIA1 and VEGF’s role as a predictive indicator of pre-term births in women living with HIV initiating ART before conception AUC 0.82 and p =0.0047

**Figure S2:**
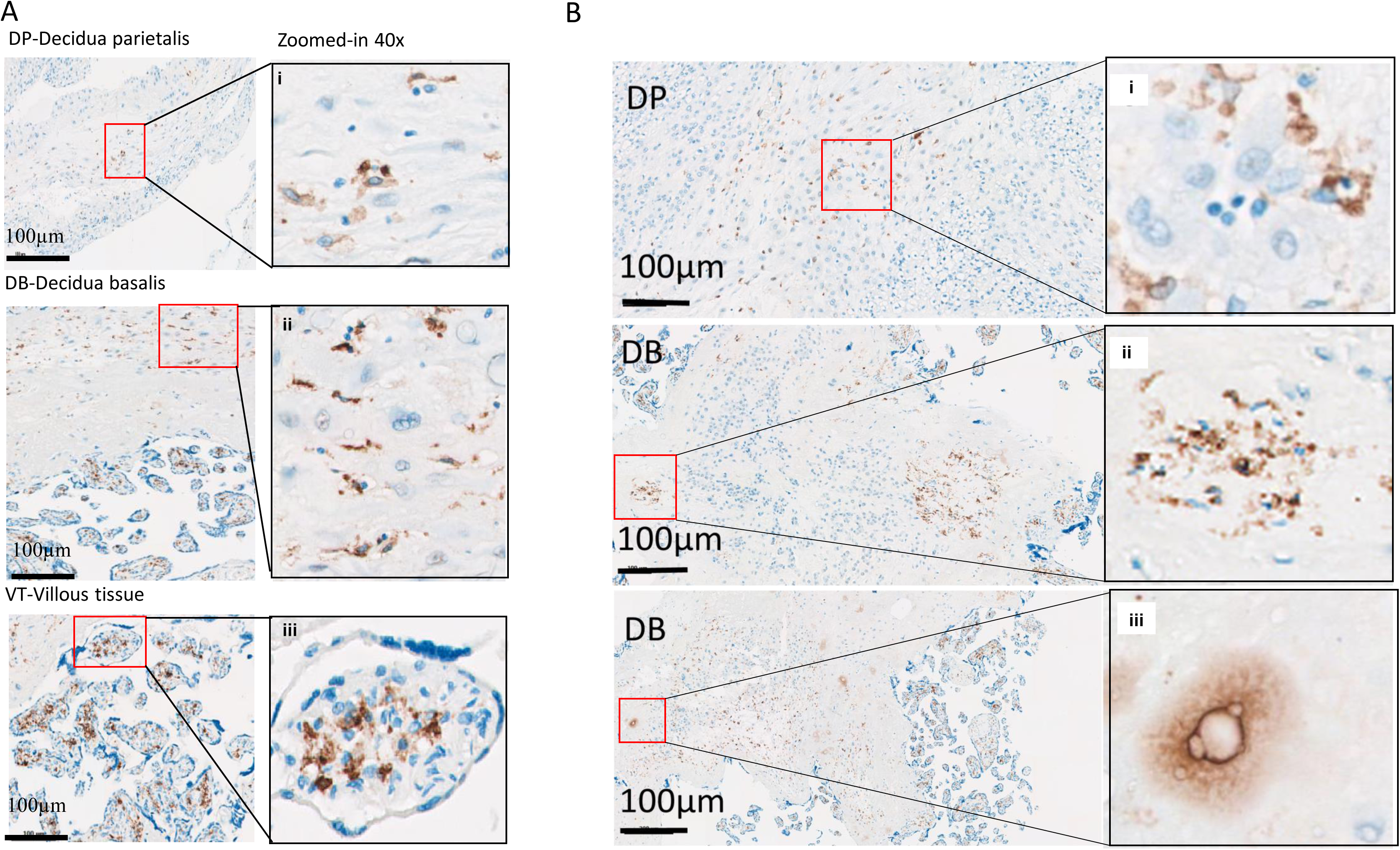
FXIIIA1 Staining in villous tissue, decidua basalis and decidua parietalis. A. Representative images of term HIV negative placental compartments showing the distribution of FXIIIA1+ cells at a scale of 100µm and the red squares zoomed-in at 40X. FXIIIA1 staining was associated with smaller nuclei in DP (i), striated in DB (ii), and enriched in VT (iii). B. In the decidua, two distinct patterns of FXIIIA1 staining are depicted at a scale of 100µm. The "striated" pattern exhibited a dimmer and stellated appearance, although the cells are less abundant and compressed. In decidua parietalis (DP), there was an association of FXIIIA1 staining with smaller nuclei (i). In some decidua basalis (DB) sections, staining revealed cells aggregating in "lumps" (ii), some of which appeared artificial but are characteristic of transglutaminases in tissue (iii).

**Figure S3:**
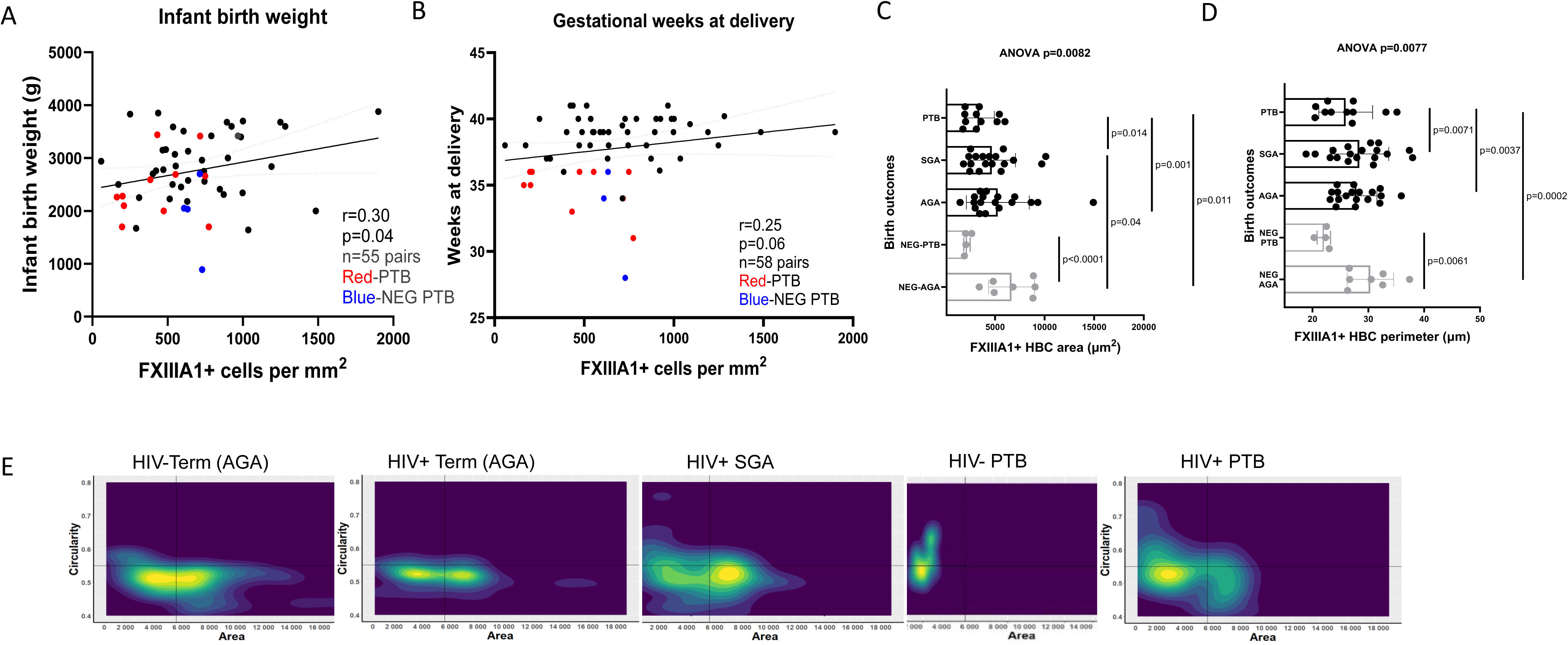
Morphological assessment of FXIIIA1+ cells and their correlation with infant weight and gestation. A. Correlation between FXIIIA1 expression and infant birthweight, with a significant association observed (p=0.04). Preterm births (PTB) are indicated in red for PPLH and in blue for uninfected mothers B. Correlation between FXIIIA1 expression and gestational weeks at delivery, with a marginal association observed (p=0.06). Preterm births (PTB) are indicated in red for PPLH and in blue for uninfected mothers Quantitation of morphological feature of area ANOVA p=0.0082 (C) and perimeter ANOVA p=0.0077 C. (D) occupied by Hofbauer cells. D. A 2D density plot showing the relationship between area and circularity (E) in various birth outcomes. Each dot represents mean ± SD of 5 regions of interest from each placenta analysed. Lines and bars represent the mean and standard deviation, respectively. The overall repeated measures ANOVA p value is shown as well as the the Two-tailed P values from Mann-Whitney U test is shown.

**Figure S4:**
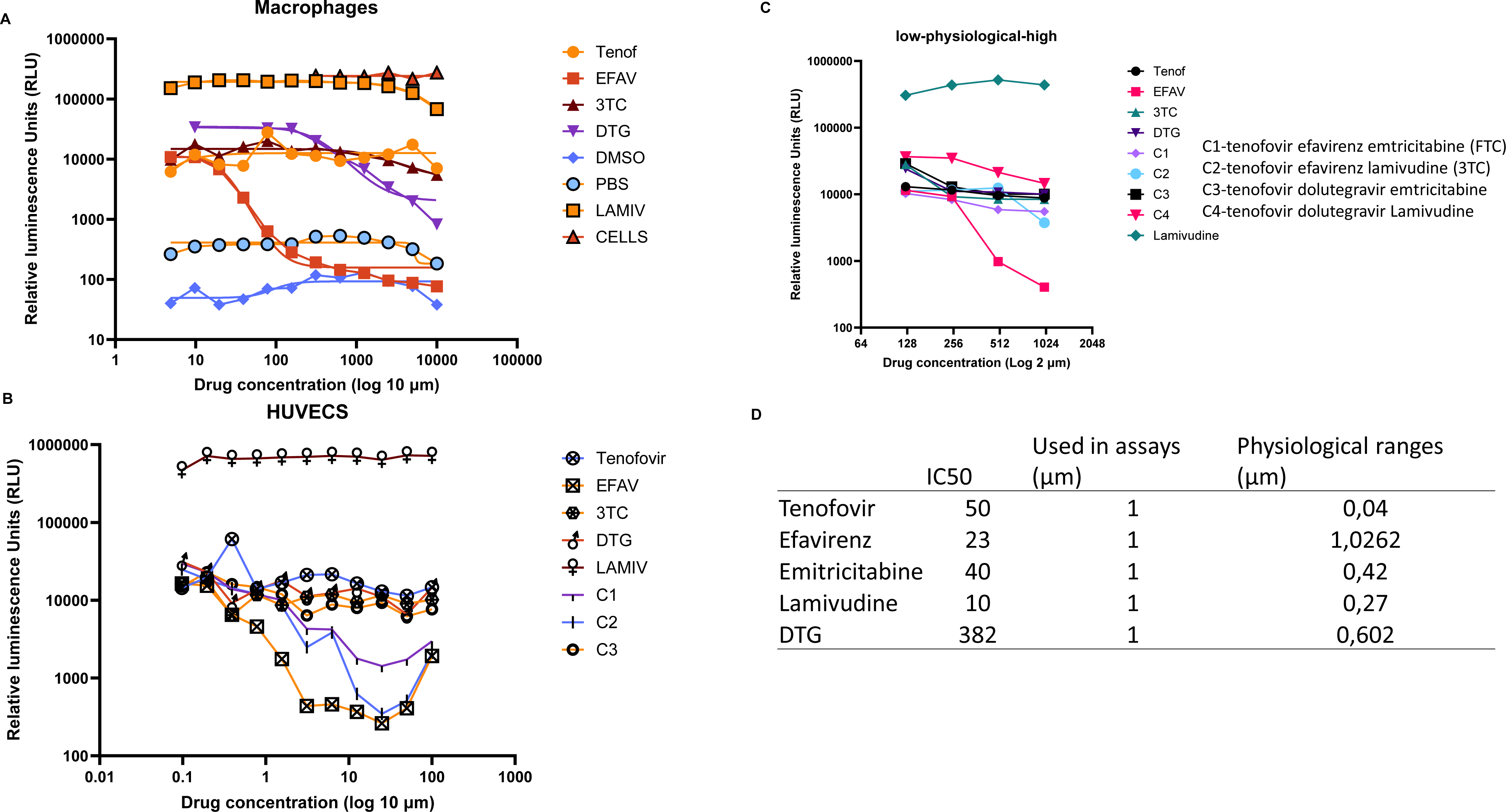
ART Cytotoxicity Measurement in Macrophages and HUVECs. Macrophages and HUVECs were treated with increasing concentrations of ART drugs, including tenofovir, emtricitabine, and lamivudine, to assess their viability. Cell viability was measured using a standard cytotoxicity assay, and the results are shown as dose-response curves. The viability of cells was plotted relative to untreated controls, indicating the non-toxic concentrations of ART used in further experiments. These findings helped determine the appropriate dosing for subsequent assays. A. A cell-viability curve shows the cytotoxicity of antiretroviral therapy (ART) drugs in macrophages and B. (B) in HUVECs at log 10 scale. C. The granular relationship is shown in HUVECs in (C) at log 2 scale. D. D shows the IC50 values obtained, the physiological ranges and concentrations used in cultures

**Table S1:**
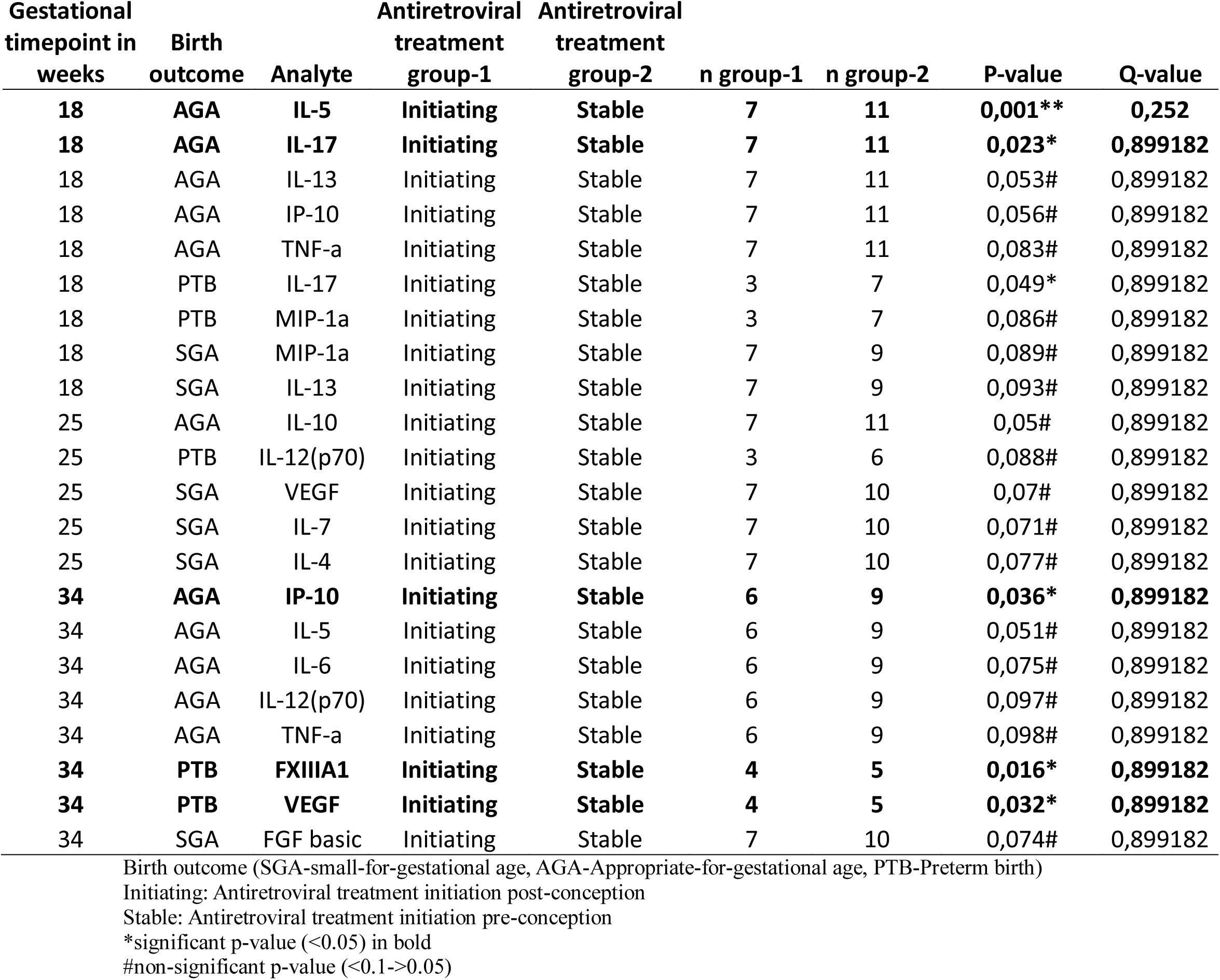
Cytokines differentially expressed between initiating and stable across different groups (birth outcomes) at the different gestational time points (18, 25 and 34)

| Gestational<br>timepoint in<br>weeks | Birth<br>outcome | Analyte | Antiretroviral<br>treatment<br>group-1 | Antiretroviral<br>treatment<br>group-2 | n group-1 | n group-2 | P-value | Q-value |
| --- | --- | --- | --- | --- | --- | --- | --- | --- |
| **18** | **AGA** | **IL-5** | **Initiating** | **Stable** | **7** | **11** | **0,001\*\*** | **0,252** |
| **18** | **AGA** | **IL-17** | **Initiating** | **Stable** | **7** | **11** | **0,023\*** | **0,899182** |
| 18 | AGA | IL-13 | Initiating | Stable | 7 | 11 | 0,053# | 0,899182 |
| 18 | AGA | IP-10 | Initiating | Stable | 7 | 11 | 0,056# | 0,899182 |
| 18 | AGA | TNF-a | Initiating | Stable | 7 | 11 | 0,083# | 0,899182 |
| 18 | PTB | IL-17 | Initiating | Stable | 3 | 7 | 0,049* | 0,899182 |
| 18 | PTB | MIP-1a | Initiating | Stable | 3 | 7 | 0,086# | 0,899182 |
| 18 | SGA | MIP-1a | Initiating | Stable | 7 | 9 | 0,089# | 0,899182 |
| 18 | SGA | IL-13 | Initiating | Stable | 7 | 9 | 0,093# | 0,899182 |
| 25 | AGA | IL-10 | Initiating | Stable | 7 | 11 | 0,05# | 0,899182 |
| 25 | PTB | IL-12(p70) | Initiating | Stable | 3 | 6 | 0,088# | 0,899182 |
| 25 | SGA | VEGF | Initiating | Stable | 7 | 10 | 0,07# | 0,899182 |
| 25 | SGA | IL-7 | Initiating | Stable | 7 | 10 | 0,071# | 0,899182 |
| 25 | SGA | IL-4 | Initiating | Stable | 7 | 10 | 0,077# | 0,899182 |
| **34** | **AGA** | **IP-10** | **Initiating** | **Stable** | **6** | **9** | **0,036\*** | **0,899182** |
| 34 | AGA | IL-5 | Initiating | Stable | 6 | 9 | 0,051# | 0,899182 |
| 34 | AGA | IL-6 | Initiating | Stable | 6 | 9 | 0,075# | 0,899182 |
| 34 | AGA | IL-12(p70) | Initiating | Stable | 6 | 9 | 0,097# | 0,899182 |
| 34 | AGA | TNF-a | Initiating | Stable | 6 | 9 | 0,098# | 0,899182 |
| **34** | **PTB** | **FXIII A1** | **Initiating** | **Stable** | **4** | **5** | **0,016\*** | **0,899182** |
| **34** | **PTB** | **VEGF** | **Initiating** | **Stable** | **4** | **5** | **0,032\*** | **0,899182** |
| 34 | SGA | FGF basic | Initiating | Stable | 7 | 10 | 0,074# | 0,899182 |

Birth outcome (SGA-small-for-gestational age, AGA-Appropriate-for-gestational age, PTB-Preterm birth)
Initiating: Antiretroviral treatment initiation post-conception
Stable: Antiretroviral treatment initiation pre-conception
\*significant p-value (<0.05) in bold

**Graphical abstract (optional).**
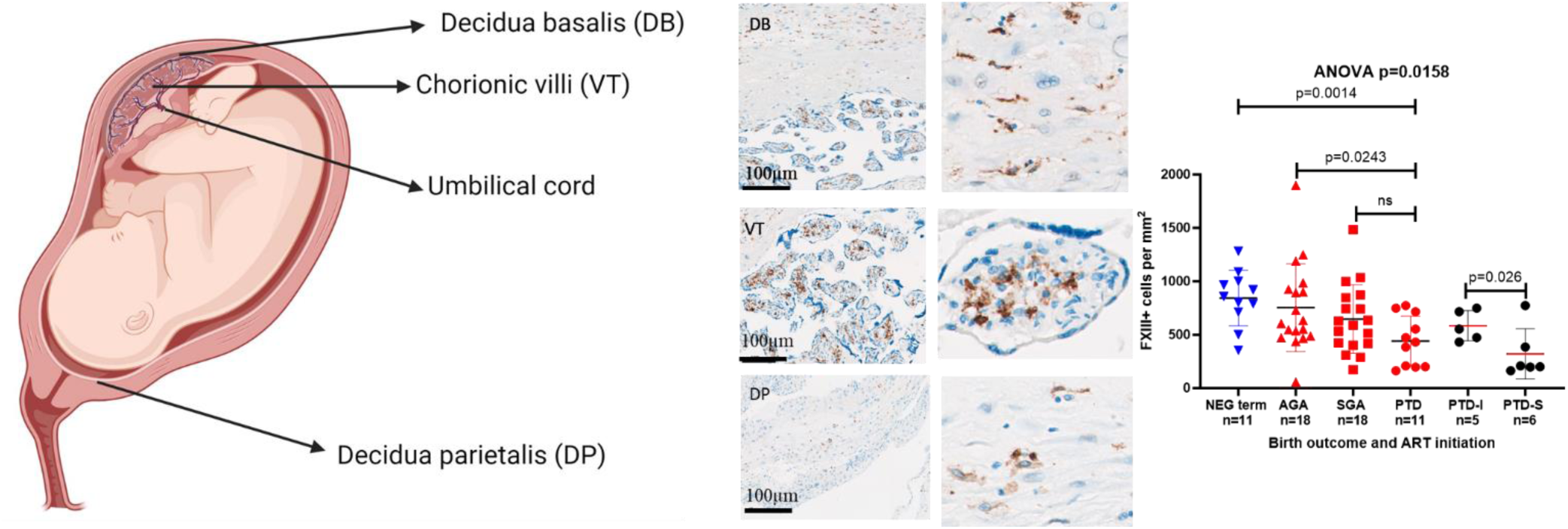
Our data suggest that low FXIIIA1 expression on Hofbauer cells is implicated in preterm delivery and from PPLH we noted reduced FXIIIA1+cells in placenta from women who initiated ART before (stable group) and not post conception (initiating group).

